# The impact of similarity metrics on cell type clustering in highly multiplexed in situ imaging cytometry data

**DOI:** 10.1101/2023.01.18.524659

**Authors:** Elijah Willie, Pengyi Yang, Ellis Patrick

## Abstract

Highly multiplexed *in situ* imaging cytometry assays have enabled researchers to scrutinize cellular systems at an unprecedented level. With the capability of these assays to simultaneously profile the spatial distribution and molecular features of many cells, unsuper-vised machine learning, and in particular clustering algorithms, have become indispensable for identifying cell types and subsets based on these molecular features. However, the most widely used clustering approaches applied to these novel technologies were developed for cell suspension technologies and may not be optimal for *in situ* imaging assays. In this work, we systematically evaluated the performance of various similarity metrics used to quantify the similarity between cells when clustering. Our results demonstrate that performance in cell clustering varies significantly when different similarity metrics were used. Lastly, we propose FuseSOM, an ensemble clustering algorithm employing hierarchical multi-view learning of similarity metrics and self-organizing maps (SOM). Using a stratified subsam-pling analysis framework, FuseSOM exhibits superior clustering performance compared to the current best-practice clustering approaches for *in situ* imaging cytometry data analysis.

## 1 Introduction

Technological advancements over the past decade have provided researchers the capability to simultaneously measure multiple molecular features in tissue at subcellular resolution [1]. Key technologies that are pioneering a new era for spatially-resolved proteomics include Imaging Mass Cytometry (IMC) [2], Multiplexed Ion Beam Imaging by Time Of Flight (MIBI-TOF) [3], Co-Detection by indEXing (CODEX) [4] and its successor Phenocycler. These technologies can measure approximately 50-100 features with high throughput, enabling researchers to address complex questions about the spatial distribution and interaction of various types of cells *in situ* [5]. Before interrogating the spatial relationships between cells however, a ubiquitous analytical step when analyzing highly multiplexed imaging data is defining functionally distinct cell types.

Unsupervised clustering algorithms are valuable tools for discovering both known and novel cell types in highly multiplexed data, even in cases where prior knowledge of the cell types present in an experiment is lacking [6]. In our review of the literature, over 70 percent of manuscripts employing highly multiplexed imaging data for analysis utilized one of three clustering algorithms. IMC and MIBI-TOF data was predominately clustered using either Phenograph [7], a graph-based Louvain community detection method, or FlowSOM [8], a self-organising map approach, while CODEX data was predominately clustered using X-shift, a KNN algorithm accessible through the Vortex GUI[9]. In the remaining manuscripts, other Louvain and Leiden graph-based community detection algorithms, hierarchical clustering and K-means clustering were used.

Despite their popularity in imaging modalities, Phenograph, FlowSOM and X-shift were developed in 2015 and 2016 for suspension cytometry technologies which do not share all of the same technical limitations and noise profiles with tissue based imaging technologies. Additionally, in practice these methods are often employed to generate a large set of candidate clusters which using expert domain knowledge are then manually clustered, refined and annotated based on biological features such as key marker expression, cell localization. Following this, there exist multiple avenues for further exploration of how clustering algorithms could be tailored for multiplexed imaging data.

Choosing an appropriate similarity metric is crucial for clustering algorithms as it de-termines how points in a dataset, in our case cells, are partitioned into clusters. Different similarity metrics, often referred to as distance metrics, can yield different clusters. While the Euclidean distance is commonly used in many clustering algorithms, recent studies have shown that correlation-based metrics such as Pearson or Spearman correlation perform better when clustering in other multiplexed technologies [10, 11]. Evaluating the performance of different similarity metrics for defining cell types in multiplexed imaging data may guide the improvement or development of new clustering algorithms which are optimal for these exciting technologies.

In this study, we systematically assess the performance of various distance-and correlation-based metrics across 15 imaging datasets, using multiple performance metrics. We also compare the performance of best-practice clustering methods that currently employ different similarity metrics. Based on our assessment, which highlights the benefits of combining information from multiple similarity metrics, we introduce a new clustering algorithm called FuseSOM. FuseSOM utilizes self-organizing maps (SOM) and combines multiple similarity metrics through multi-view ensemble learning and hierarchical clustering. This algorithm aims to accurately and robustly identify cell types in multiplexed in situ imaging cytometry assays. Overall, our work demonstrates the impact of similarity metrics on clustering cells in multiplexed imaging cytometry data and proposes FuseSOM as a promising method for the analysis of such data.

## Results

### Evaluating the impact of similarity metrics

To assess the impact of similarity metrics on clustering performance, we performed hierarchical clustering on a MIBI-TOF dataset using either correlation-based metrics (Pearson, Spearman, and Cosine) or distance-based metrics (Euclidean, Manhattan, and Maximum) [12]. To assess performance, we compared the clusters from the hierarchical clustering to the manually curated cell type labels identified in the manuscript (**Figure 1**). On average, correlation-based metrics outperform distance-based metrics by 8.0% for ARI, 10.7% for NMI, 1.70% for FM-index, and 8.0% for F-Measure. Furthermore, there is evidence that clustering performance is improved when using correlation-based metrics compared to distance-based metrics, as seen across three evaluation metrics.

**Figure 1:**
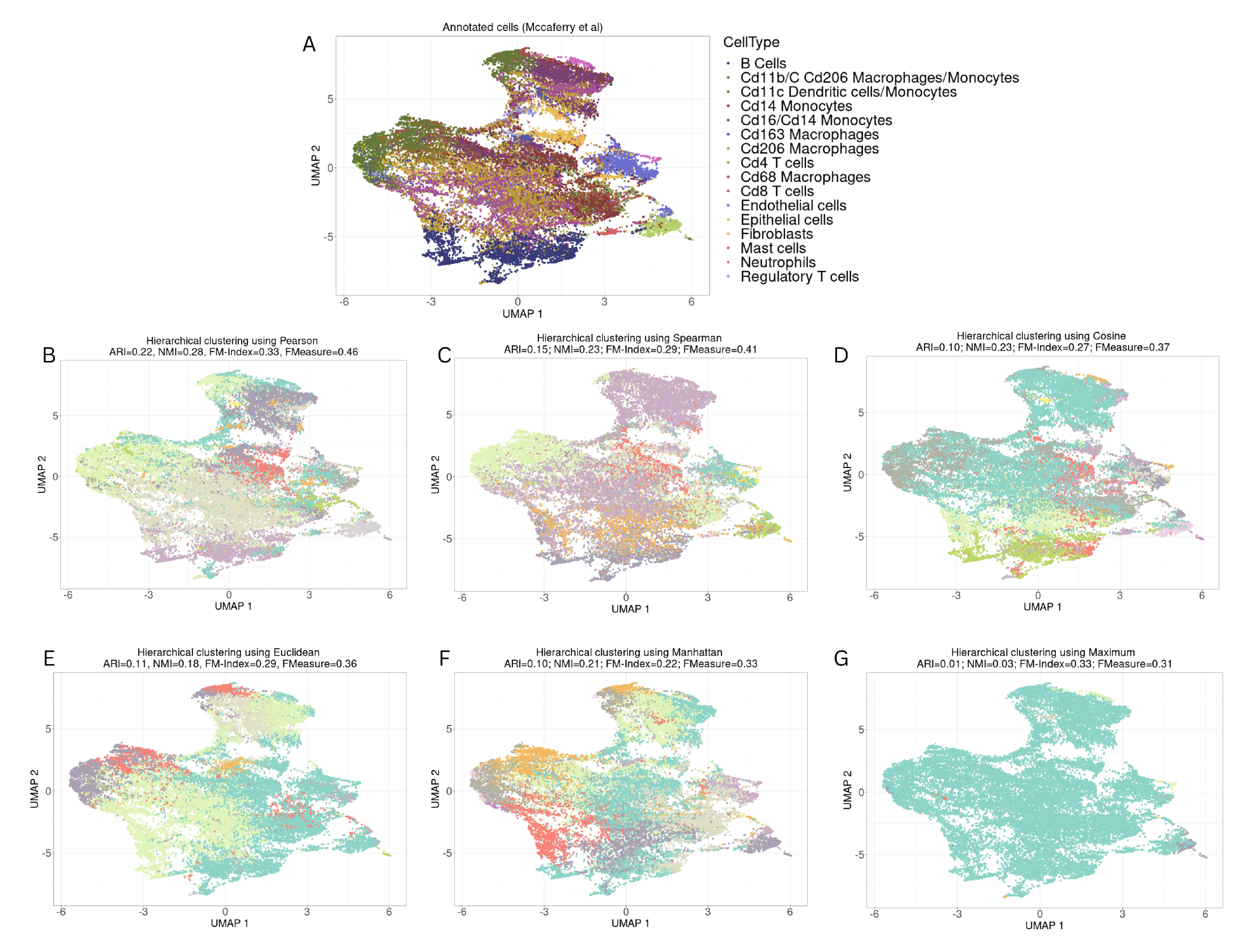
UMAP visualization of cells from a sample imaging dataset. (A) Cells colored by annotations from original study [12]. (B) Hierarchical clustering using Pearson’s Correlation and concordance quantified by ARI, NMI, FM-Index, and FMeasure. (C) Hierarchical clustering using Cosine’s Distance (D) Hierarchical clustering using Spearman’s Correlation. (E) Hierarchical clustering using Euclidean distance. (F) Hierarchical clustering using Manhattan distance. (G) Hierarchical clustering using Maximum distance.

To provide a comprehensive assessment of the performance of the similarity metrics, we quantified the clustering performance of the metrics on 15 multiplexed in situ imag-ing cytometry datasets (**Table 1**). These datasets were chosen as each had some manual intervention when cell type labels were defined. Each dataset was randomly subsampled to 20K cells five times, and each subset was clustered using hierarchical clustering with all the similarity metrics. Finally, the average was taken across the five subsets. Across all 15 datasets, correlation-based metrics consistently outperformed distance-based metrics (**Figure 2**), more accurately recapitulating the manually curated cell-type labels from their original publications. These results show the efficacy of correlation-based metrics in hierar-chical clustering.

**Figure 2:**
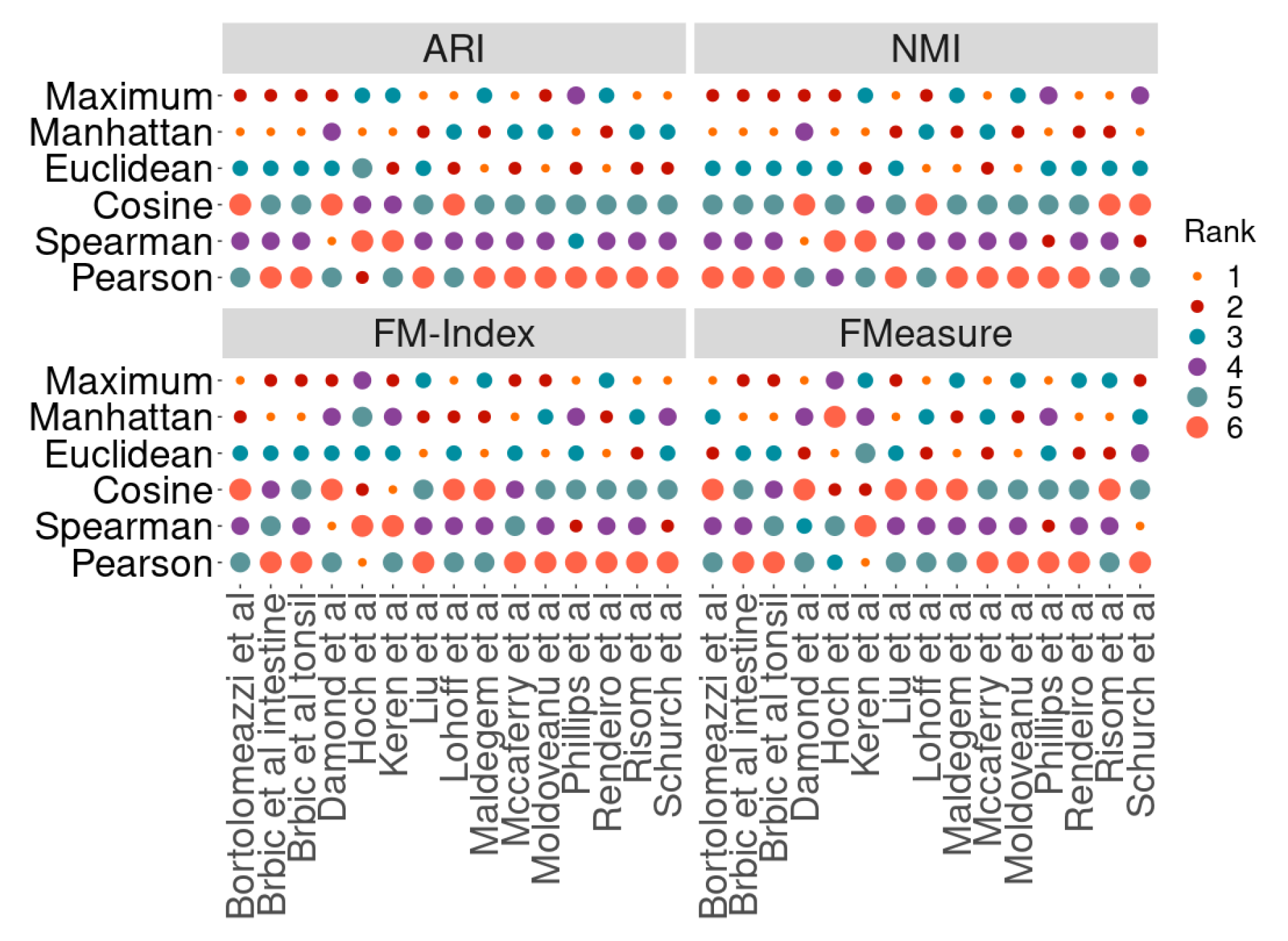
Benchmarking similarity metrics on agglomerative clustering of 15 multiplexed imag-ing datasets. Each dataset was subsetted to 20K cells five times, and the average clustering score was recorded. Results were ranked in descending order across all similarity metrics and datasets by each evaluation metric. A larger circle size indicates better performance. Correlation-based metrics are consistently ranked higher than distance-based metrics across most datasets.

**Table 1:** Imaging Datasets Used

| Dataset | Technology | Num Markers | Num Cells | Num Celltypes | Disease | Tissue |
| --- | --- | --- | --- | --- | --- | --- |
| Shchurch et al [41] | CODEX | 49 | 258,385 | 29 | Colorectal Cancer | Colon |
| Phillips et al [42] | CODEX | 52 | 117,170 | 21 | Lymphoma | Skin |
| Brbic et al [43] | CODEX | 48 | 248,285 | 21 | None | Small intestine/colon |
| Brbic et al [43] | CODEX | 44 | 219,926 | 13 | Normal/Bartlett's esophagus | Tonsil |
| Moldoveanu et al [44] | IMC | 12 | 227,592 | 10 | Melanoma | Skin |
| Van Maldegem et al [45] | IMC | 17 | 282, 837 | 16 | Lung Cancer | Lung |
| Hoch et al [14] | IMC | 41 | 864,263 | 10 | Melanoma | Skin |
| Rendeiro et al [46] | IMC | 38 | 515,791 | 17 | Covid-19 | Lung |
| Damond et al [47] | IMC | 36 | 252,059 | 16 | Type 1 Diabetes | Pancreas |
| Bortolomeazzi et al [48] | IMC | 30 | 218,615 | 9 | Colorectal Cancer | Colon |
| Risom et al [49] | MIBI-TOF | 22 | 69,672 | 23 | Breast Cancer | Breast |
| Keren et al [3] | MIBI-TOF | 16 | 201,656 | 6 | TN Breast Cancer | Various |
| McCafferry et al [12] | MIBI-TOF | 37 | 30,943 | 16 | Tuberculosis | Various |
| Liu et al [50] | MIBI-TOF | 12 | 345,490 | 8 | Various Cancers | Various |
| Lohoff et al [51] | seqFISH | 50 | 57,536 | 24 | None | Mouse embryos |

Next, we assessed if the performance differences between correlation and Euclidean distance are consistent across multiple clustering methods. We reconfigured PhenoGraph, FlowSOM, and K-means clustering to use correlation instead of euclidean [7, 8]. As previously, the 15 datasets were randomly subsetted to 20K cells five times, and after clustering, the average score was taken across all evaluation metrics. While there does not appear to be a significant benefit in clustering performance for PhenoGraph, the overall PhenoGraph performance for both distance measures is worse when compared to the correlation-based distances (**Figure 3**) and (**Supplementary Figure S1**). For K-means, hierarchical clustering, and FlowSOM, we observe Pearson correlation performing better than Euclidean distance across all evaluation metrics. See (**Figure 3**) and (**Supplementary Figure S1**).

**Figure 3:**
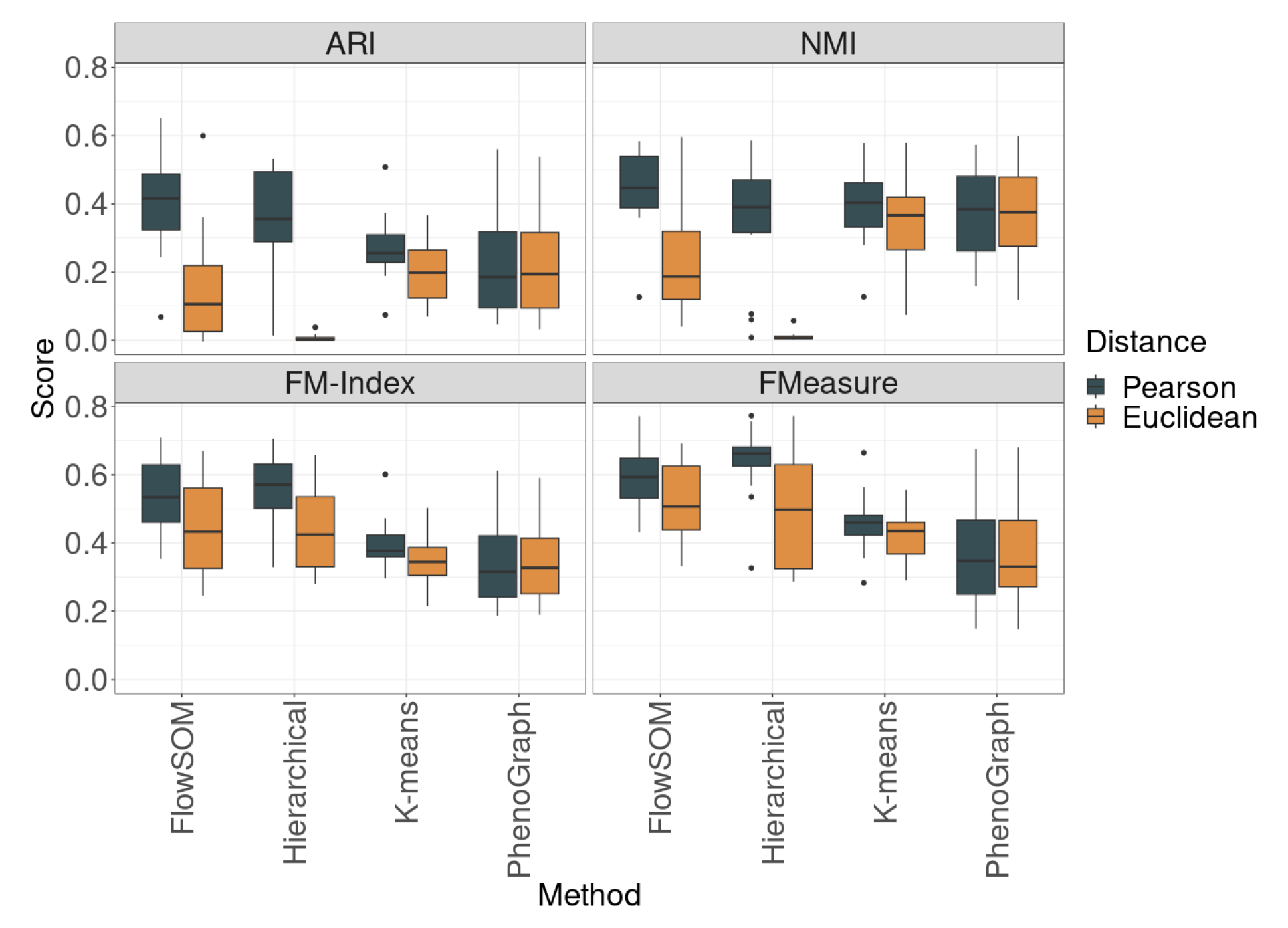
Boxplots of clustering performance of four clustering methods using Pearson correlation and euclidean across four evaluation metrics (ARI, NMI, FM-Index, and FMeasure). For three of the four methods, Pearson on average performs better than euclidean for clustering. This is not evident for PhenoGraph.

### Combining similarity metrics is beneficial

Given the performance differences between the similarity metrics, we next wondered if combining multiple metrics using strategies such as multi-view ensemble learning would further improve performance [13]. To evaluate the efficacy of combining multiple distance metrics for clustering, we performed a comparison study combining various combinations of Pearson, Spearman, Cosine, and the Euclidean distance. First, all possible combinations of metrics were used to cluster the prototypes generated by the SOM algorithm. Of all the combinations, combining Pearson, Spearman, Cosine, and Euclidean consistently provided the best score across all four evaluation metrics (**Figure 4**). The combination of Pearson, Spearman, and Cosine also provided strong results, which imply that the addition of Euclidean does not markedly improve overall clustering performance.

**Figure 4:**
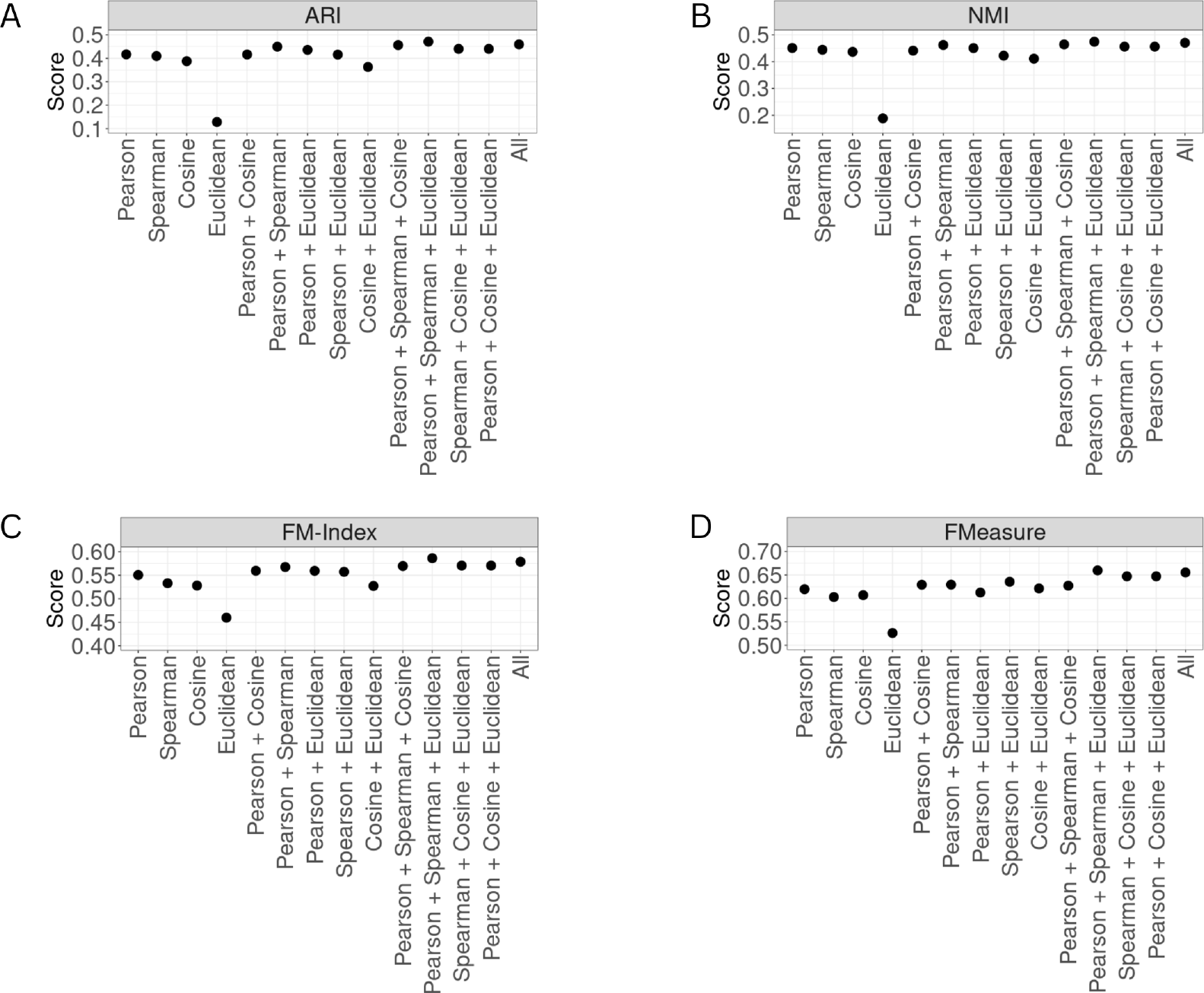
Scatter plots of average clustering performance of all combinations of Pearson, Cosine, Spearman, and Euclidean. Across all performance metrics, there is evidence that combining all distances provides a better signal for clustering.

### FuseSOM combines self-organizing maps with multi-view ensemble learning of similarity metrics

Here, we introduce FuseSOM for the clustering of highly multiplexed imaging data. FuseSOM leverages all the ideas already discussed by combining similarity metrics with a Self Organizing Map and multiview hierarchical clustering to define cell types robustly (**Figure 5**). In comparison to FlowSOM which uses Euclidean distance by default, FuseSOM has superior performance in our stratified subsampling analysis framework which demonstrates that multiview ensemble of similarity metrics does provide a more robust clustering (**Figure 6**). The performance gain is particularly evident when looking at ARI and NMI, with average differences in scores being 32% for ARI, 27% for NMI, 10% for FM-Index, and 9.0% for FMeasure.

**Figure 5:**
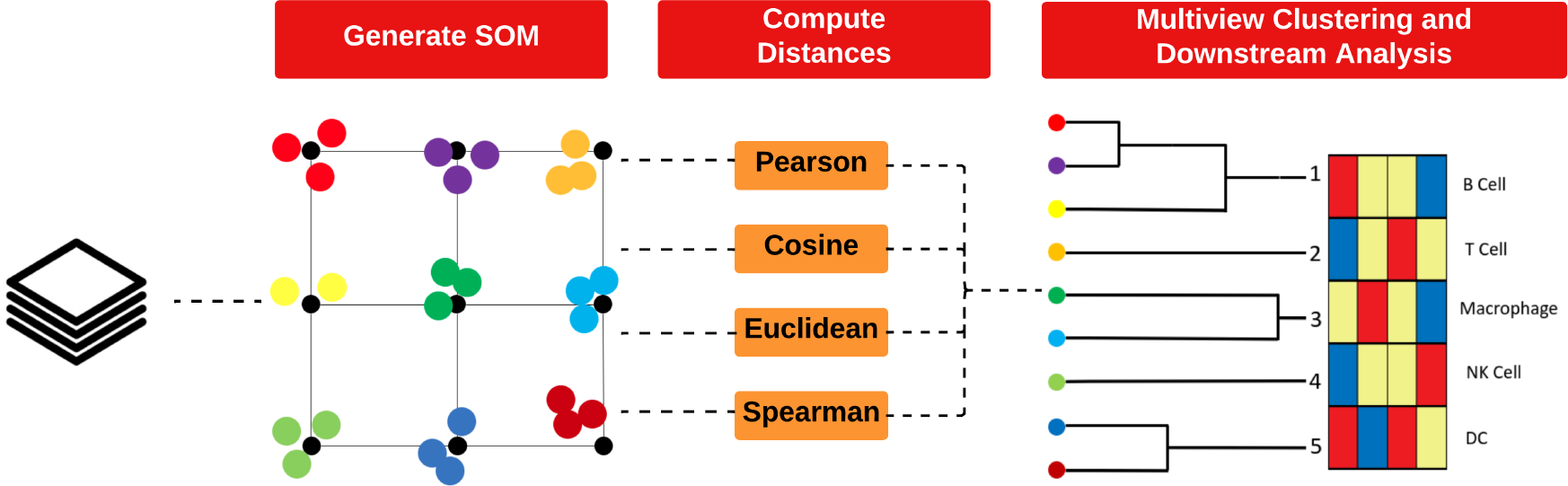
Overview of FuseSOM: This new scalable algorithm uses (i) A Self Organizing Map to reduce the dimension of the data while preserving its topological structure; (ii) A multiview integration of various similarity metrics to capture all relevant signals; and (iii) Hierarchical clustering to generate a clustering solution for further downstream analysis.

**Figure 6:**
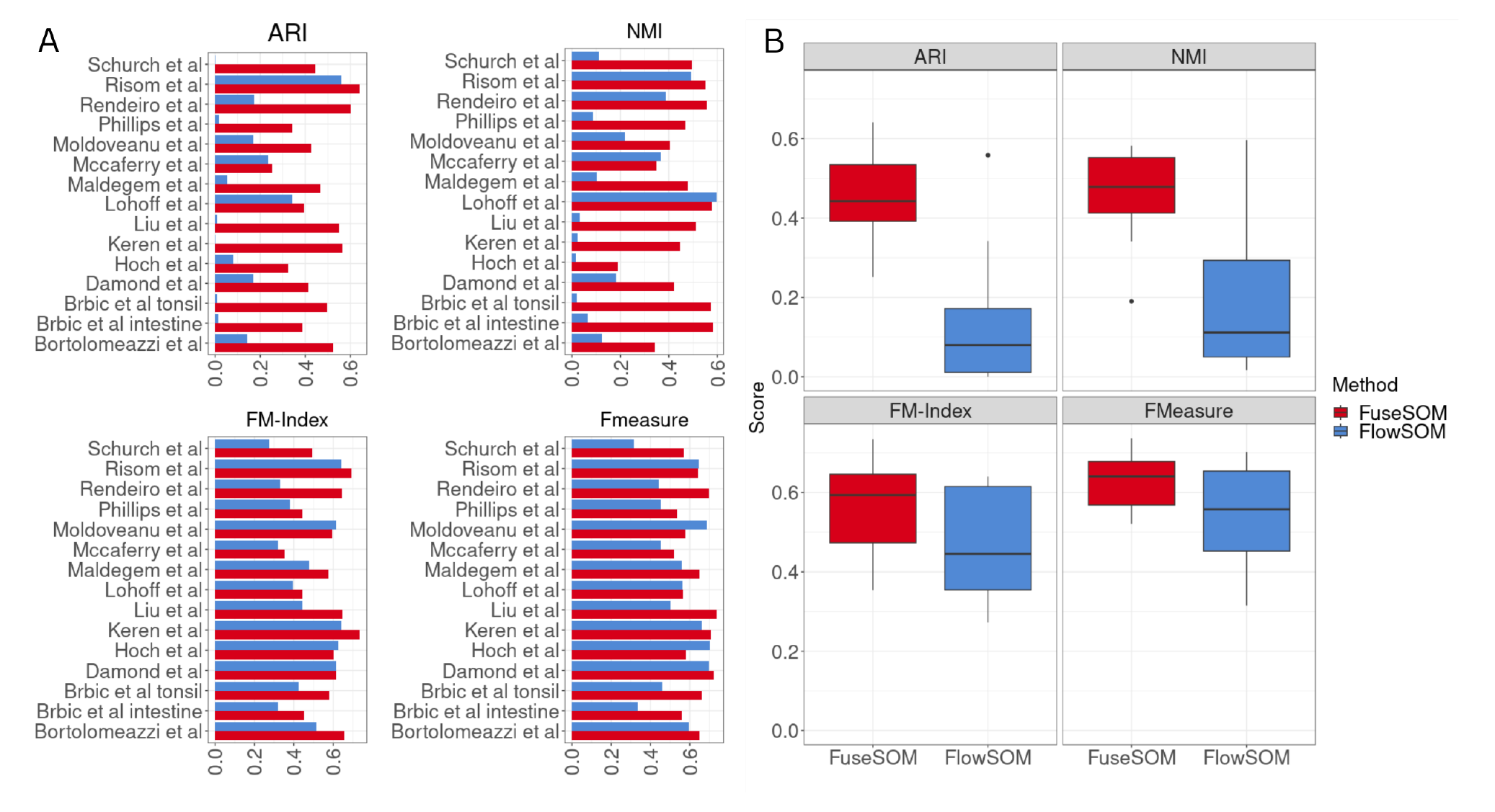
(A) Bar plots of average clustering performance for FuseSOM and FlowSOM showed for all 15 datasets across four performance metrics (ARI, NMI, FM-Index, and FMeasure). For a majority of the datasets, FuseSOM outperforms FlowSOM. (B) Boxplots for the average cluster-ing performance across all datasets. For all four evaluation metrics, FuseSOM shows an overall higher average score.

Several methods for estimating the number of clusters have been implemented in the FuseSOM R package. The relative error (RE) between the predicted number of clusters and the number of clusters used in the corresponding manuscripts was used to assess the accuracy of the cluster estimation methods. The Jump and Discriminant method tends to overestimate the actual number of clusters while the others tend to underestimate the actual number of clusters (**Figure** 7). Next, FuseSOM was used to cluster the cells in the datasets using the number of clusters estimated by each method. For both ARI and NMI, the Jump and Discriminant methods appear to have the highest average score across all datasets (**Figure 7**). This demonstrates that the Discriminant and Jump methods have the greatest average accuracy in predicting the number of clusters chosen in the original manuscripts. This is further reflected in the clustering performance based on the predicted number of clusters.

**Figure 7:**
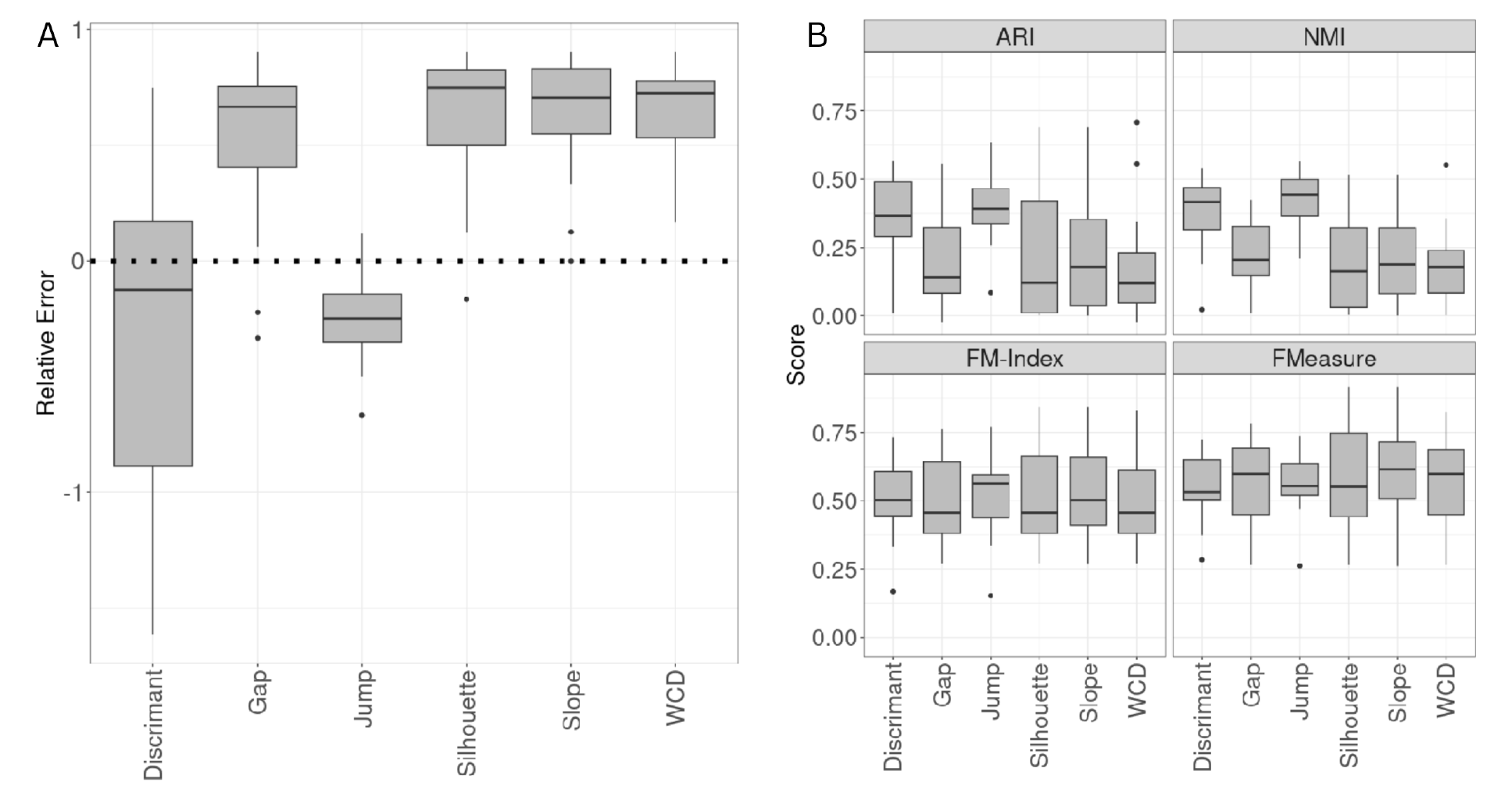
(A) Bar plots of cluster estimation performance for methods included in FuseSOM across all 15 datasets. The relative error metric was used to gauge performance. The Jump and Discriminant methods are the top performers. The dashed line represents a relative error of zero. In contrast, values above this line indicate an underestimation of the true number of clusters, and values below indicate an overestimation of the true number of clusters. (B) Boxplots of clustering performance based on the estimated number of clusters for methods included in FuseSOM. The Jump and Discriminant methods are top performers across all evaluation metrics.

To investigate the running time and memory usage of FuseSOM, we applied FuseSOM to the Hoch dataset [14]. This dataset contains 800K cells across 41 markers. Next, clustering was performed in increments of 100K cells (8 clustering solutions in total) to gauge how memory and running are affected by an increasing number of cells. Clustering was performed on an 11th Gen Intel® Core™ i7-1165G7 @ 2.80GHz with four cores and 32 GB of memory. Memory usage scales linearly with the size of the data (**Supplmentary Figure S2a**). Doubling the size of the data requires twice as much memory. FuseSOM can process a dataset with 800K cells using only 8GB of memory, implying that FuseSOM can scale to datasets with more than a million cells with moderate memory usage. FuseSOM is also efficient, being able to process 800K cells in under 5 minutes (**Supplementary Figure S2b**). The computational time scales sub-linearly as the size of the dataset increases. For example, the running time is not doubled by doubling the data size.

## Discussion

In this work, we performed a comparative analysis of the performance of various similarity metrics for clustering highly multiplexed *in situ* imaging cytometry assays. Using multiple clustering methods across multiple similarity metrics, we demonstrated that the choice of similarity metric does affect the clustering performance of highly multiplexed cytometry *in situ* imaging data with correlation-based metrics on average out-performing distance-based metrics. We then leveraged these findings to develop a novel multi-view clustering algorithm called FuseSOM and demonstrated its ability to recover semi-supervised cell-type annotations across various datasets from differing imaging technologies with reasonable accuracy. Our results comprehensively demonstrate the impact of similarity metric choice on cell type clustering in highly multiplexed imaging cytometry data, and highlight the need to develop new best-practice clustering algorithms for these technologies.

While we have demonstrated that correlation metrics are often superior to distance metrics for multiplexed imaging data, we have not demonstrated why. We do however demonstrate that this phenomena is consistent across the imaging platforms. Our hypothesis is that distance-based metrics such as Euclidean and Manhattan are sensitive to the scaling of the data and therefore are susceptible to changes in the expression of markers across images or even different regions of the tissue imaged. However, correlation-based metrics such as Pearson and Spearman are scale-invariant and, therefore, could be less susceptible to changes in the expression of markers driven by technical artifacts. As correlation-based metrics only consider the relative expression between markers, we suspect this makes them more robust and therefore are more accurate in capturing cell type specific expression trends in highly multiplexed in situ imaging cytometry data.

For many clustering algorithms, the number of clusters is a parameter that must be chosen before executing the algorithm. Some algorithms, including graph and density based, can estimate the number of clusters as part of the algorithm, however, even for these methods other parameters indirectly affect the number of clusters. For example, the size of the neighborhood in density-based methods can affect the number of clusters. Selecting a suitable number of clusters should always be viewed in the context of the application and while it can be guided by quantitative approaches, like those we have implemented in FuseSOM, should ultimately be assess qualitatively. For example, to identify rare cell types in biological data, one might need to deeply cluster the data to find smaller populations of cells. Or, after over-clustering data to identify smaller cell subsets, expert domain knowledge may be used to reduce the number of clusters so that the resulting clusters were biologically sensi-ble or interpretable. We expect that clustering will always benefit from manual intervention and scrutiny to decide the number of appropriate clusters for a dataset and we advocate for leveraging prior domain knowledge when annotating cell types.

## Methods

### Imaging datasets collection and processing

To benchmark the performance of FuseSOM on imaging datasets from various technologies, we selected a set of 15 datasets generated using different imaging technologies. In addition, we selected datasets with human intervention in manually gating cell populations or merging biologically similar clusters. Datasets were sourced from major databases, including Zenodo, Figshare, and Mendeley. When available, we used the version data that had been processed as described in the original manuscript. We also used the same markers for clustering as described in the manuscript. The imaging technologies used included Co-detection by indexing(CODEX) (four datasets), Imaging Mass Cytometry(IMC) (six datasets), Multiplexed ion beam imaging by the time of flight (MIBI-TOF) (four datasets), and sequential Fluorescence In Situ Hybridization (seqFISH) (one dataset) [3, 4, 15, 16]. Next, we employed a substratification framework for each dataset to evaluate clustering methods. This framework takes an imaging dataset as input, and five stratified samples are returned. Sub-stratification is done to account for possible variability in clustering results. This framework takes as input and returns five stratified datasets. Stratification is done by selecting 80% of cells from each annotated class [10].

### Similarity metrics

Six types of metrics across two classes that are predominantly used across machine learning clustering literature were used in this study. The two classes include correlation-based and distance-based. The distance-based metrics were Euclidean, Manhattan, and Maximum distance, while the correlation-based metrics included Pearson correlation, Spearman correlation, and Cosine similarity. More formally, let *x_im_* and *x_jm_* denote the expression of a marker *m* = 1*, …, M* in cell *i* = 1*, …, N* and cell *j* = 1, …, *N*, where *G* and *N* are the total number of markers and cells, respectively. Led *D* = *d_ij_* be a distance matrix where *d_ij_* represents the distance between *cell_i_* and *cell_j_*. We can then define the distance-based metrics as follows:

Euclidean distance,

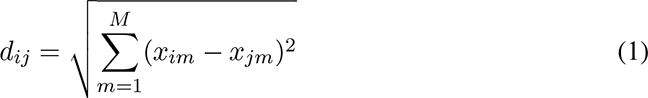

Manhattan distance,

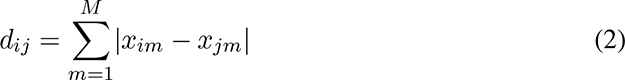

Maximum distance,

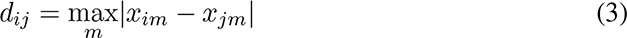

Similarly, the correlation-based metrics can be defined as follows:

Pearson distance,

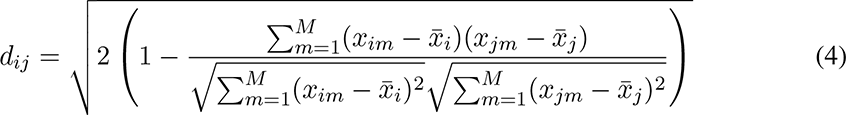

Spearman distance,

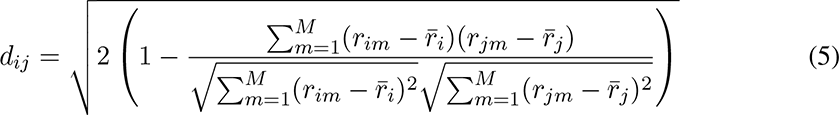

Cosine distance

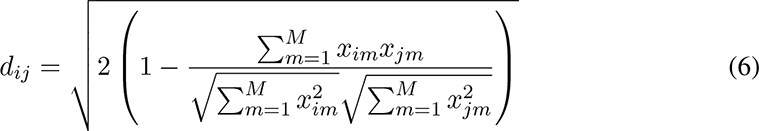

where *r_ij_* is the rank of marker *m* in *cell_i_*, 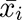 is the mean expression of *cell_i_*, 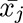 is the mean expression of *cell_j_*, 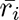 is the mean expression rank of *cell_i_*, and 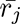 is the mean expression rank of *cell_j_*,

### The hierarchical clustering and self-organizing map algorithms

We used the agglomerative hierarchical clustering algorithm with varying similarity metrics to investigate the effect of similarity metrics on clustering algorithms. Agglomerative clustering is a type of clustering where each data point starts as its cluster. At every step, two clusters are merged using a linkage function. This step repeats until a single cluster comprises the entire dataset. There are many options for a linkage function. We opted to use the average linkage function, which merges two clusters based on the average distance between pairs of points between two clusters [17]. This algorithm outputs a tree-like structure called a dendrogram which can be cut at any level to generate *k* clusters. We also used the FlowSOM algorithm proposed by [8]. This algorithm uses a Self Organizing Map (SOM) architecture to generate prototypes which are then clustered using hierarchical consensus clustering using euclidean distances to generate meta-clusters. These meta-clusters are then projected back on the original dataset to obtain clusters.

### FuseSOM

Here the FuseSOM algorithm is described. The algorithm starts by generating a Self Organizing Map. A Self Organizing Map is a type of dimensionality reduction algorithm that maps points in a high dimensional space (*d >* 2) to a lower dimensional space (*d* = 2). The SOM architecture preserves the topological relationships between points when reduced to a lower dimension. The SOM also provides a set of points called prototypes which are representations of points in the higher dimensional space. Like cluster centers in the k-means algorithm, many points can be mapped to a single prototype. For a more thorough treatment of SOMs, see [18]. The *YASOMI* package was obtained and modified to implement the self-organizing map used in FuseSOM [19].

In this work, we generate a SOM, and the prototypes are used for clustering. After clustering, the clusters are projected back to the original data to classify the original data points. The SOM algorithm requires a 2-d grid(x,y) size, which determines the number of prototypes. Grid sizes of varying shapes are allowed. However, square grids are typically used. To estimate the size of the grid for a dataset, we use the method described in [20]. This method computes the number of eigenvalues of a covariance matrix significantly different from the Tracy-Widom distribution [21].

Next, multiview integration combines the Pearson correlation distance, Cosine distance, Spearman correlation, and Euclidean distance between the prototypes to generate a final distance matrix for clustering. We adopted a multiview integration method to combine the four transformed matrices [22]. The *analogue* package is used to perform the multiview integration [23]. The *psych* package transforms the similarity matrices into distance matrices [24]. Finally, to generate final cluster labels using the integrated distance matrices, hierarchical clustering using the average linkage function is used. The *FCPS* package was used for hierarchical clustering [25].

### Estimating the number of clusters

For most clustering algorithms, the number of clusters *k* is an important hyperparameter that must be set. To this end, many methods have been developed to help practitioners choose an appropriate number for their dataset. We have included well-known methods for estimating the number of clusters as part of the FuseSOM package. These methods include the Gap statistic, the Slope statistic, the Jump statistic, the Silhouette statistic, and the within-cluster distance (WCD) [26–29]. We also developed a method for estimating the number of clusters based on maximum clusterability discriminant projection pursuit. To accomplish this, we couple hierarchical clustering with discriminant analysis and multimodality testing to estimate the number of clusters [30–35]. First, we generate a dendrogram using hierarchical clustering with average linkage. Next, for each node in the resulting tree, we project the two classes onto a line such that both classes are well separated. See (**Supplementary Figure S3**). The Dip test for multimodality testing is then applied to the distribution of the points along this line [34]. See (**Figure 7**). The family-wise error rate (FWER) is controlled using the method described in [36]. Finally, the number of nodes with significant p-values is returned as the number of clusters.

### Evaluation metrics

To evaluate clustering performance for clustering solutions generated across methods, we used a set of methods; the Adjusted Rand Index (ARI), Normalized Mutual Information (NMI), Fowlkes-Mallows index (FM-Index), and the F-Measure [37–40]. All these metrics take values between 0 and 1, with 0 being no similarity and 1 being perfect similarity.

## Data availability

Publicly available data were used for all evaluations. All data were downloaded as described in the originating manuscripts.

## Code availability

The *FuseSOM* R package is available on Bioconductor and is available under the GPL-3 license. All code for analysis done can be found at Github

## Acknowledgments

The authors thank all their colleagues, particularly at The University of Sydney, Sydney Precision Data Science Centre and Charles Perkins Centre for their support and intellectual engagement.

The following sources of funding for each author are gratefully acknowledged: the AIR@innoHK programme of the Innovation and Technology Commission of Hong Kong to EP and PY. Australian Research Council Discovery Early Career Researcher Award (DE200100944) funded by the Australian Government to EP; National Health and Medical Research Council (NHMRC) Investigator Grant (1173469) to P.Y.; and the University of Sydney Postgraduate Excellence Award for E.W.; The funding sources had no impact on the study design; in the collection, analysis, and interpretation of data, the writing of the manuscript, and in the decision to submit the manuscript for publication.

## Author contributions

EP and EW conceived and designed the study. EP and EW led the method development and guided the evaluation data analysis with input from PY. EW curated the imaging data, implemented all data analytics, and developed the corresponding R code. All authors wrote, read, reviewed the manuscript, and approved the final version.

## Supplementary Figures

**Supplementary Figure S1:**
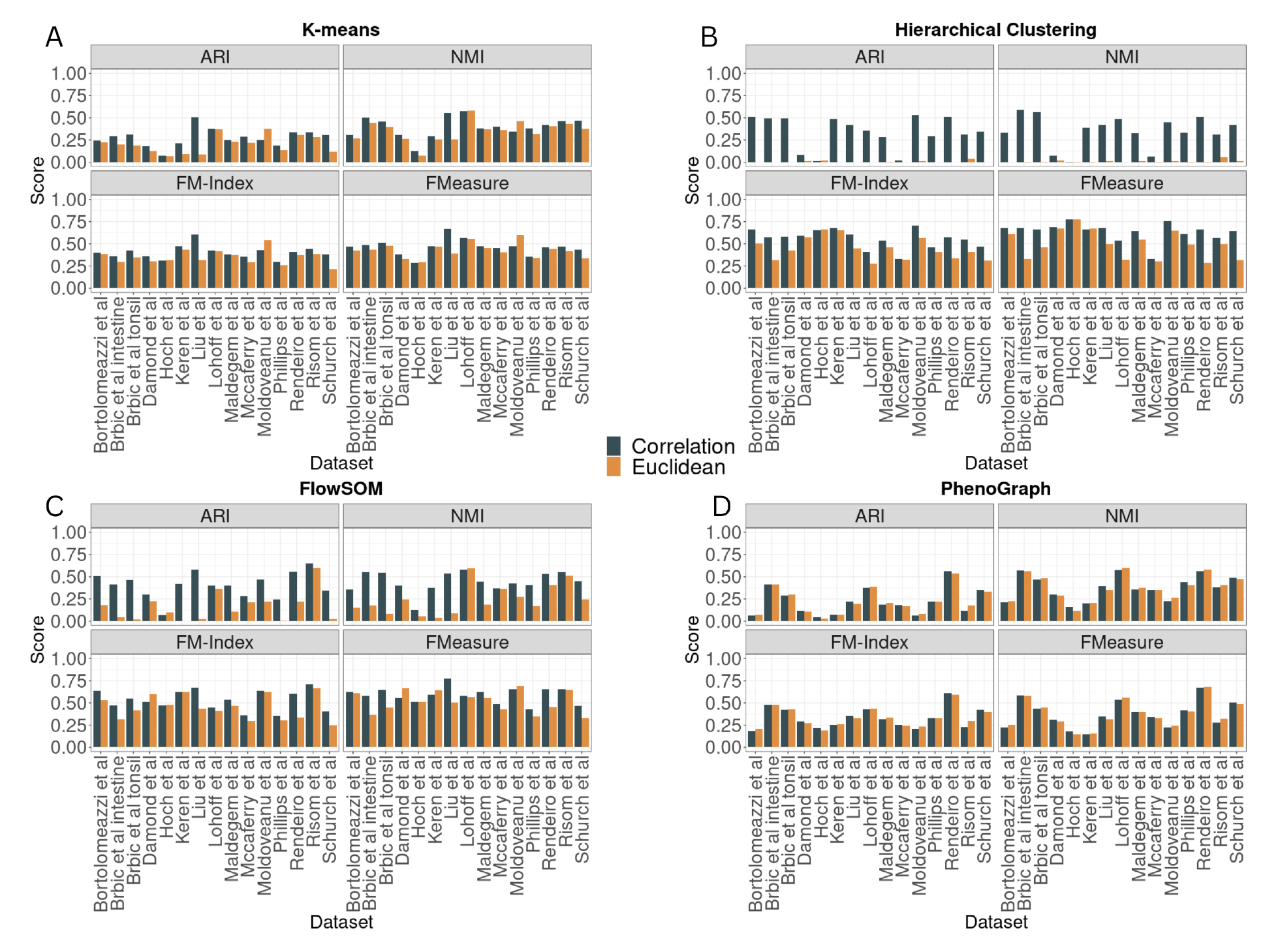
(A) Bar plots of K-means clustering of a subset of 20K cells for 15 datasets using Pearson correlation and Euclidean distance. K-means with Pearson correlation, on average, performs better than Euclidean for most of the datasets across all evaluation metrics. (B) Bar plots of Hierarchical clustering of a subset of 20K cells for 15 datasets using Pearson correlation and Euclidean distance. On average, hierarchical clustering with Pearson correlation performs better than Euclidean for most datasets across all evaluation metrics. C) Bar plots of FlowSOM clustering of a subset of 20K cells for 15 datasets using Pearson correlation and Euclidean distance. FlowSOM clustering with Pearson correlation, on average, performs better than Euclidean for most datasets across all evaluation metrics. D) Bar plots of PhenoGraph clustering of a subset of 20K cells for 15 datasets using Pearson correlation and Euclidean distance. There is no evidence of performance differences between Pearson and Euclidean.

**Supplementary Figure S2:**
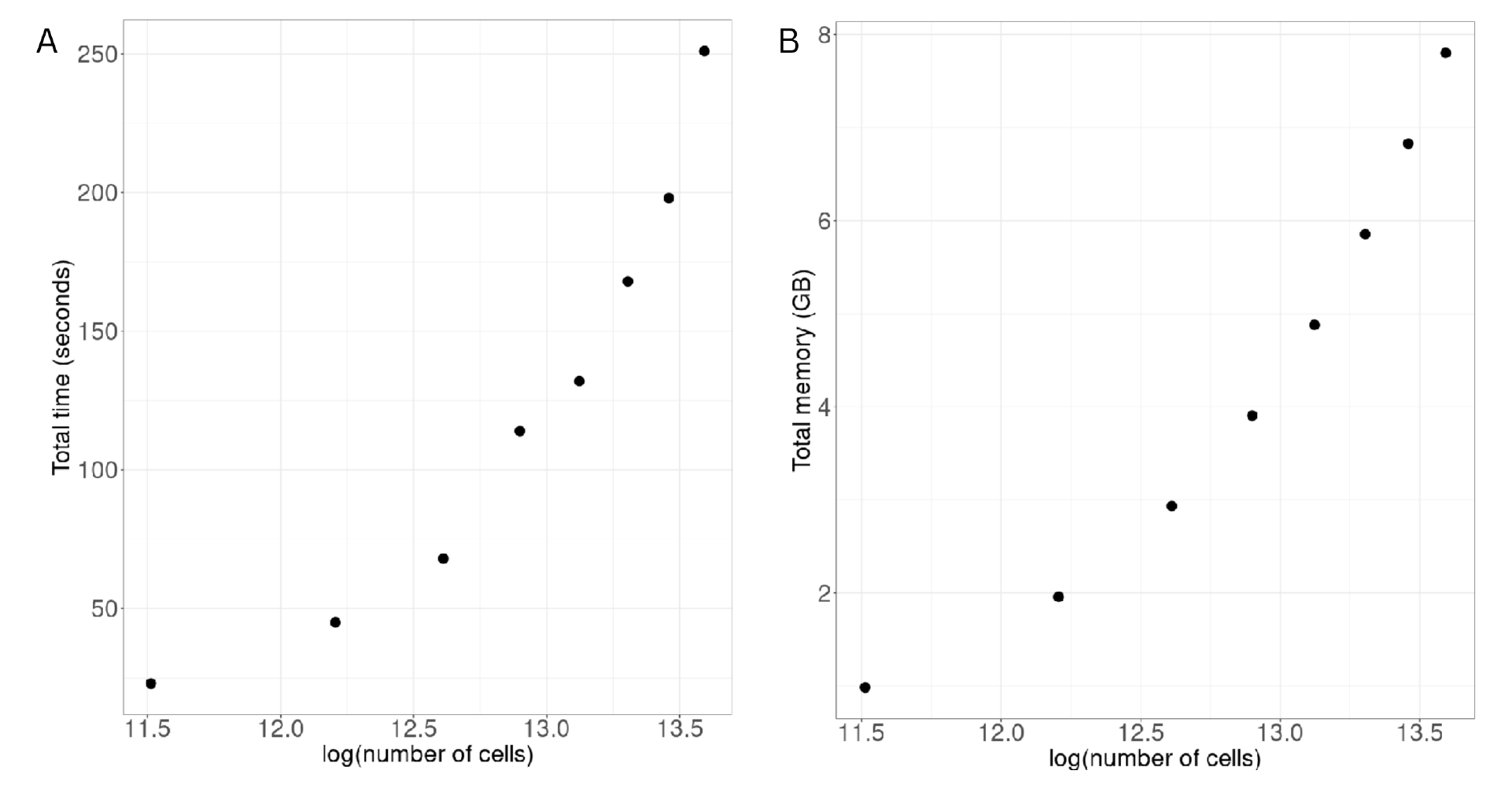
(A) Scatter plot of total memory usage for FuseSOM across an increasing number of cells. (B) Scatter plot of total running time for FuseSOM across an increasing number of cells.

**Supplementary Figure S3:**
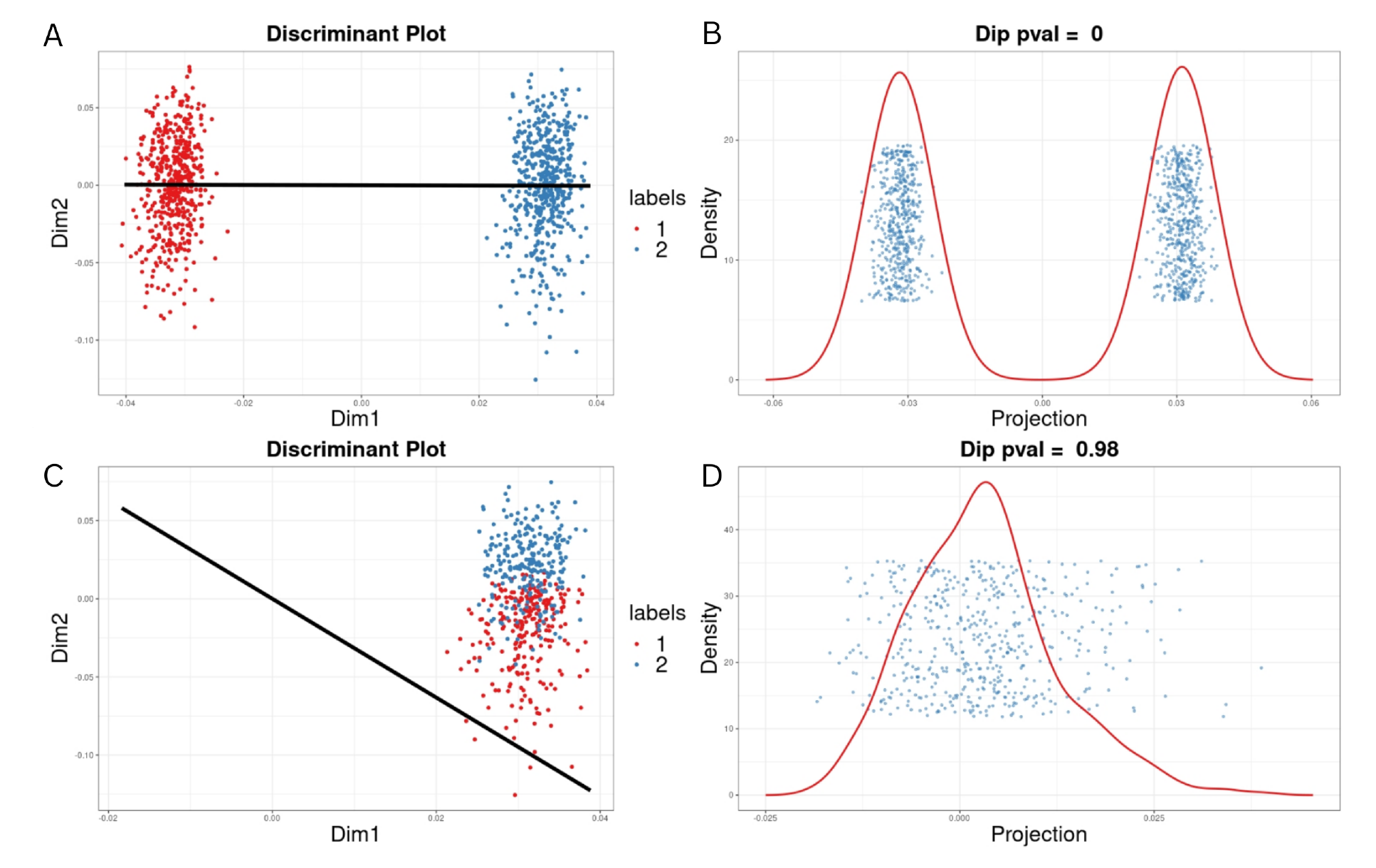
An example of discriminant coordinates cluster number estimation. (A) and (C) Shows two classes separated by a discriminant line in black. (B) and (D) Shows the Dip test applied to the projection of the two classes onto this discriminant line. Note that for (B), a significant p-value is obtained, reflecting the class separation. For (D), the opposite is observed. P-values < 2*e*^−16^ were rounded to 0.

